# RecA structure kinetics determine SOS dynamics and predict spontaneous cell death

**DOI:** 10.64898/2026.10.19.760072

**Authors:** Joseph J. Pollacco, Thomas Haygarth, Agathe Couturier, Christian Lesterlin, David J. Sherratt, Ruth E. Baker, Philip K. Maini, Stephan Uphoff

## Abstract

Even under optimal growth conditions, bacterial cells experience endogenous stresses that threaten cellular function. Individual cells sporadically activate stress responses to restore homeostasis, but some cells ultimately die. The SOS response to DNA damage is particularly heterogeneous, but the origin of its spontaneous activation and its relationship to cell fate remain unclear. Here, we imaged fluorescently labelled RecA in single *Escherichia coli* cells under steady-state unperturbed growth using microfluidics. We find that RecA* structures assemble frequently near replication forks and precede SOS response pulses. RecA* structure lifetime quantitatively encodes the magnitude of SOS induction: longer-lived structures promote greater LexA degradation and larger SOS pulses. Thus, cell-to-cell variability in spontaneous SOS activation reflects differences in the repair kinetics of endogenous DNA damage events. By tracking cell lineages over time, we further show that RecA* structures are strong and early predictors of cell fate. Elevated RecA* structure presence emerges up to six generations before spontaneous cell death in over 75% of cases, before other physiological disturbances become detectable. Together, our results establish spontaneous SOS activation as a quantitative readout of endogenous DNA damage processing and identify failure to resolve such damage as a major route to spontaneous bacterial cell death.

## Introduction

During favourable conditions, bacteria tightly control core cellular processes to maintain homeostasis and maximise growth. In contrast, processes related to cellular stress tolerance display substantial variability (Silander et al., 2012). Such heterogeneity may arise from imperfect regulation, but it may also reflect deliberate adaptive responses to intrinsic fluctuations or stress. Even in the absence of external perturbations, cells must continuously cope with natural decay of lipids, proteins, and DNA (Lindahl, 1993) as well as self-generated internal stresses from reactive metabolic byproducts (Burcham, 1999; Linster et al., 2013; Imlay, 2019). To manage these threats, cells activate stress responses to restore homeostasis and repair damage, but these mechanisms often impose significant energetic penalties and physiological burdens, such as growth arrest, altered metabolism, or mutagenesis (Vincent and Uphoff, 2020). As a result, cells must balance the benefits of stress protection against the cost of reduced proliferation. Heterogeneous activation of stress responses within isogenic bacterial populations has been proposed as a strategy to manage this trade-off (Ackermann, 2015).

Faithful DNA replication is essential for bacterial proliferation, requiring the complete duplication of the chromosome prior to cell division (Cox et al., 2000). However, replication fork progression is continually challenged by endogenous DNA damage, including abasic sites, oxidative lesions, single-stranded gaps, and double-strand breaks, as well as obstacles such as DNA-bound proteins (Burcham, 1999; Xia et al., 2019). The bacterial SOS response is a broadly conserved gene regulatory mechanism for detecting and repairing DNA damage, extensively characterized in the model species *Escherichia coli* (Radman, 1975; Little and Mount, 1982; Janion, 2008). DNA breaks or gaps are processed to generate single-stranded DNA, onto which RecA is loaded to form nucleoprotein filaments (RecA*) (Bell and Kowalczykowski, 2016). RecA* filaments serve two separate functions, performing homologous recombination repair and triggering autocleavage of the transcriptional repressor LexA. Degradation of LexA relieves repression of the SOS regulon (Giese et al., 2008; Cory et al., 2024). In *E. coli*, this regulon comprises over 30 genes involved in DNA repair and damage tolerance (Courcelle et al., 2001). Beyond DNA repair, SOS influences broader stress-associated phenotypes including cell filamentation, mutagenesis, virulence, horizontal gene transfer, and phage defence (Baharoglu and Mazel, 2014; Kelley, 2006).

Stress responses, including the SOS response, have been widely studied in bacteria under acute perturbations, such as antibiotic treatment or environmental shocks, where cells experience rapid and severe damage. In these contexts, the mechanisms and functional benefits of stress responses are more readily identifiable. However, such conditions differ fundamentally from steady-state growth, where damage arises sporadically and asynchronously across individual cells, and may accumulate gradually (Yang et al., 2023). Indeed, single-cell studies have revealed that a small fraction of cells spontaneously generate transient pulses of SOS expression (Livny and Friedman, 2004; Pennington and Rosenberg, 2007; Massoni et al., 2012; Helfrich et al., 2015; Turnbull et al., 2016; Mavridou et al., 2018). The magnitude and duration of SOS pulses are heterogeneous (McCool et al., 2004; Kamenšek et al., 2010; Mrak et al., 2007; Jones and Uphoff, 2021; Jaramillo-Riveri et al., 2022; Galbusera et al., 2025). How exactly this heterogeneity arises and whether it reflects unavoidable noise in gene regulation or a precise adaptation to endogenous DNA damage remains unclear, as does its impact on cell physiology during normal growth.

Here, we address these questions by directly visualizing SOS response dynamics in single *E. coli* cells during unperturbed growth. Using a chromosomally encoded RecA-YFP fusion (Wiktor et al., 2021), we simultaneously monitor endogenous DNA damage events and SOS gene expression via time-lapse imaging of bacteria in microfluidic devices (Wang et al., 2010; Jones and Uphoff, 2021). We show that variability in the kinetics of RecA* structures explains heterogeneity in SOS induction. Remarkably, the presence of RecA* structures is an early predictor of diverse cell fates, including lysis, growth arrest, and filamentation, several generations before detectable changes in growth or morphology. The results show that heterogeneous SOS activation is a quantitatively controlled response to endogenous DNA damage, and reveal a temporal sequence of how cell death follows rare failures of otherwise robust cellular processes.

## Results

### Spontaneous RecA* structures precede SOS expression pulses in single cells

To investigate the mechanisms underlying heterogeneous SOS response induction, we performed time-lapse microscopy of live *E. coli* cells expressing a functional RecA-YFP fusion construct (Renzette et al., 2005; Lesterlin et al., 2014; Amarh et al., 2018; Wiktor et al., 2021). Using a mother machine microfluidic device (Wang et al., 2010), we continuously monitored cells during unperturbed exponential growth with a generation time of 49 min under constant inflow of fresh minimal defined growth media (Fig. 1A). The fluorescence intensity from RecA-YFP serves as a reporter for SOS response gene expression. Cells additionally expressed the fluorescent protein mKate2 as a cytoplasmic marker to facilitate automatic cell segmentation and lineage tracing. As previously shown (Uphoff, 2018; Jones and Uphoff, 2021; Wiktor et al., 2021; Jaramillo-Riveri et al., 2022; Galbusera et al., 2025), cells exhibited transient SOS expression pulses (Fig. 1B,C), leading to substantial cell-to-cell heterogeneity and a long tail in the RecA-YFP fluorescence intensity distribution across the population at any given timepoint (Fig. 1B).

**Fig. 1.**
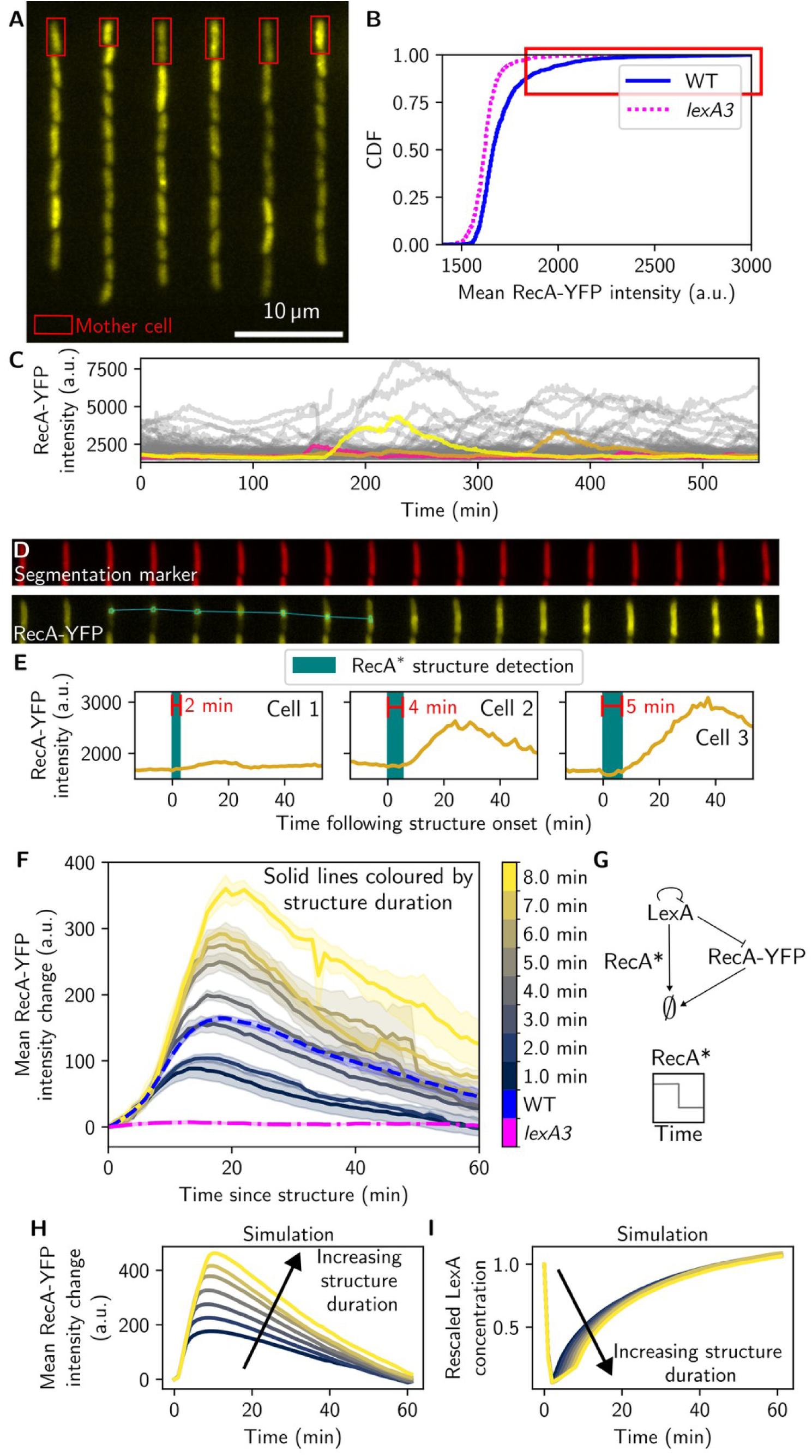
Duration of RecA* structures explains the magnitude of spontaneous SOS induction. **(A)** Mother machine snapshot showing cell-to-cell heterogeneity in RecA-YFP fluorescence (yellow). **(B)** Cumulative distribution of RecA–YFP intensity in single cells at the start of imaging for wild type (blue) and *lexA3* mutant (magenta). The high-intensity ‘SOS-induced’ subpopulation is indicated (red box). Data are pooled across experiments (WT n=4; *lexA3* n=3 biological replicates). **(C)** RecA-YFP intensity in single mother cells over time, shown for one experimental replicate. Three representative mother cells are highlighted (pink, gold, and yellow) over the rest of the population (grey). **(D)** Representative kymograph of a single mother cell (top, mKate2 segmentation marker; bottom, RecA-YFP), showing formation of a RecA* structure followed by increased RecA-YFP fluorescence. **(E)** Representative single-cell traces showing RecA-YFP intensity with detected RecA* structures for three cells. **(F)** Mean RecA-YFP fluorescence over time after RecA* structure detection for each strain (mean ± s.e.m. across biological replicates; WT n=4; lexA3 n=3). Traces for WT were split by RecA* structure duration (1–8 min, darker to lighter curves). **(G)** Schematic of a minimal model for spontaneous SOS induction: LexA represses its own expression and that of RecA-YFP, and is degraded in the presence of RecA* structures. See Supplementary information for model design and implementation. **(H)** Simulated RecA-YFP fluorescence for RecA* structures of increasing duration (1–8 min; darker to lighter curves). **(I)** Simulated LexA concentrations for RecA* structures of increasing duration (1–8 min; darker to lighter curves).

We observed localised RecA-YFP structures in approximately 5% of cells at any given time. These appeared as foci or filaments, accounting for approximately 82% and 18% of structure observations, respectively, and are hereafter jointly referred to as RecA* structures. Cells with RecA* structures had lower fluorescence intensity elsewhere in the cytoplasm, consistent with cytoplasmic RecA being recruited into dynamic nucleoprotein filaments, as shown in other works (Renzette et al., 2005; Lesterlin et al., 2014; Amarh et al., 2018; Wiktor et al., 2021; Chimthanawala et al., 2022). RecA* structures generally appeared in cells immediately prior to a transient SOS expression pulse (Fig. 1D,E). This confirms that spontaneous SOS pulses are indeed triggered by RecA* structure formation rather than by spontaneous fluctuations or noise in basal *recA* gene expression.

To ascertain the expected link between RecA* structures and SOS induction in our system, we treated cells with the antibiotic ciprofloxacin, which inhibits DNA gyrase, leading to DNA replication stalling and double-strand DNA breaks. As expected (Mok and Brynildsen, 2018; Broughton et al., 2026), the mean number of RecA* structures in cells increased quickly after treatment, followed by an increase in the mean RecA-YFP fluorescence intensity of the population (Supplementary Fig. 1A,B). We ruled out photodamage as a major cause of the observed RecA* structure events in untreated cells using control experiments at different light exposure doses (Supplementary Fig. 1C,D).

### RecA* structure kinetics explain heterogeneity in spontaneous SOS response induction

Confirming the known association between RecA* structures and SOS induction does not, however, explain why SOS pulses are so heterogeneous in magnitude. We considered if differences in the individual RecA* structures might be at the source of SOS heterogeneity. In untreated cells, cells rarely showed more than one detectable structure, so we began by classifying cells simply by structures being ‘present’ or ‘absent’. From these classifications, we produced single-cell traces of RecA-YFP fluorescence intensity, augmented with RecA* structure detections (Fig. 1E). Visual inspection suggested that the magnitude of a spontaneous SOS pulse depends on the duration of the preceding RecA* structure.

To test this quantitatively, we aligned the RecA-YFP intensity traces relative to the onset of a RecA* structure detection (Fig. 1F, blue dashed curve). This showed that RecA expression is induced immediately (within 1 min) upon RecA* structure formation, peaks by 15-20 min, and decays back to the basal expression level by 60 min. We then split and grouped the intensity traces by the duration of their associated RecA* structures and computed the mean trace for each group (Fig. 1F, coloured curves). We found that as the duration of the RecA* structure increases, so too does the peak height of RecA-YFP fluorescence intensity. Even the shortest detectable structures, lasting <1 min, resulted in a small but significant expression pulse. The total fluorescence intensity associated with a RecA* structure was also correlated with the magnitude of subsequent SOS response induction, as expected when the rate of LexA cleavage is determined by the encounter frequency with a RecA* filament (Supplementary Fig. 2). To confirm that gene expression pulses are caused by canonical SOS signalling via cleavage of LexA, we performed the same experiment and analysis in a mutant strain expressing the non-cleavable *lexA3* variant (Lin and Little, 1988). As expected, RecA* structures still formed in this mutant but did not trigger increased RecA-YFP expression (Fig. 1B,F, magenta dashed line).

These data suggest that variable durations of RecA* structures control the extent of LexA degradation, which in turn explains the variable magnitude of SOS pulses. To test this hypothesis, we devised a simple mathematical model. Guided by previous models (Aksenov, 1999; Krishna et al., 2007; Galbusera et al., 2025), we reduced the SOS response to a minimal number of components (Fig. 1G), and used ordinary differential equations (ODEs) to predict RecA and LexA concentration curves in response to RecA* structure formation (Fig. 1H,I). The curves recapitulate key features of the experimental data (Fig. 1F). Specifically, LexA is degraded at a constant rate per molecule while a RecA* structure is present, leading to an exponential decay in LexA concentration and accompanying increase in RecA expression. This gives rise to the same relationship between duration of RecA* structure and height of the RecA expression pulse as seen in experiments. Taken together, our experiments and model show that seemingly disordered spontaneous SOS responses in unperturbed cells are quantitatively controlled by the kinetics of RecA* structures. Rather than behaving as a binary on/off switch, spontaneous SOS activation is therefore finely graded according to the repair kinetics of endogenous DNA lesions.

### Genetic perturbations reveal distinct classes of RecA* repair events

Given that RecA* structure kinetics are predictive of heterogeneity in SOS response induction, we sought to examine the distribution of structure kinetics, and explore how genetic perturbations affect this distribution (Fig. 2A). As above, the *lexA3* mutant was chosen to break the feedback loop between RecA* structure formation and SOS induction. To perturb the kinetics of RecA* structures, we deleted *dinI*, which encodes a protein responsible for stabilising RecA* filaments (Lusetti et al., 2004; Renzette et al., 2007; Galkin et al., 2011). We also constructed a strain that constitutively expresses PsiB (*psiB+*), a protein normally produced by conjugative plasmids to inhibit the SOS response upon entry into a new host cell (Couturier et al., 2023). PsiB sequesters free RecA in the cell and prevents it from binding to the single stranded plasmid, thus stopping the recipient cell from costly SOS induction (Petrova et al., 2009). Although constitutive expression of PsiB in cells is arguably an artificial perturbation, it is a useful tool for our purposes, as it allows us to reduce the concentration of free RecA without interfering with *recA* gene regulation or the properties of RecA* once assembled into structures.

**Fig. 2.**
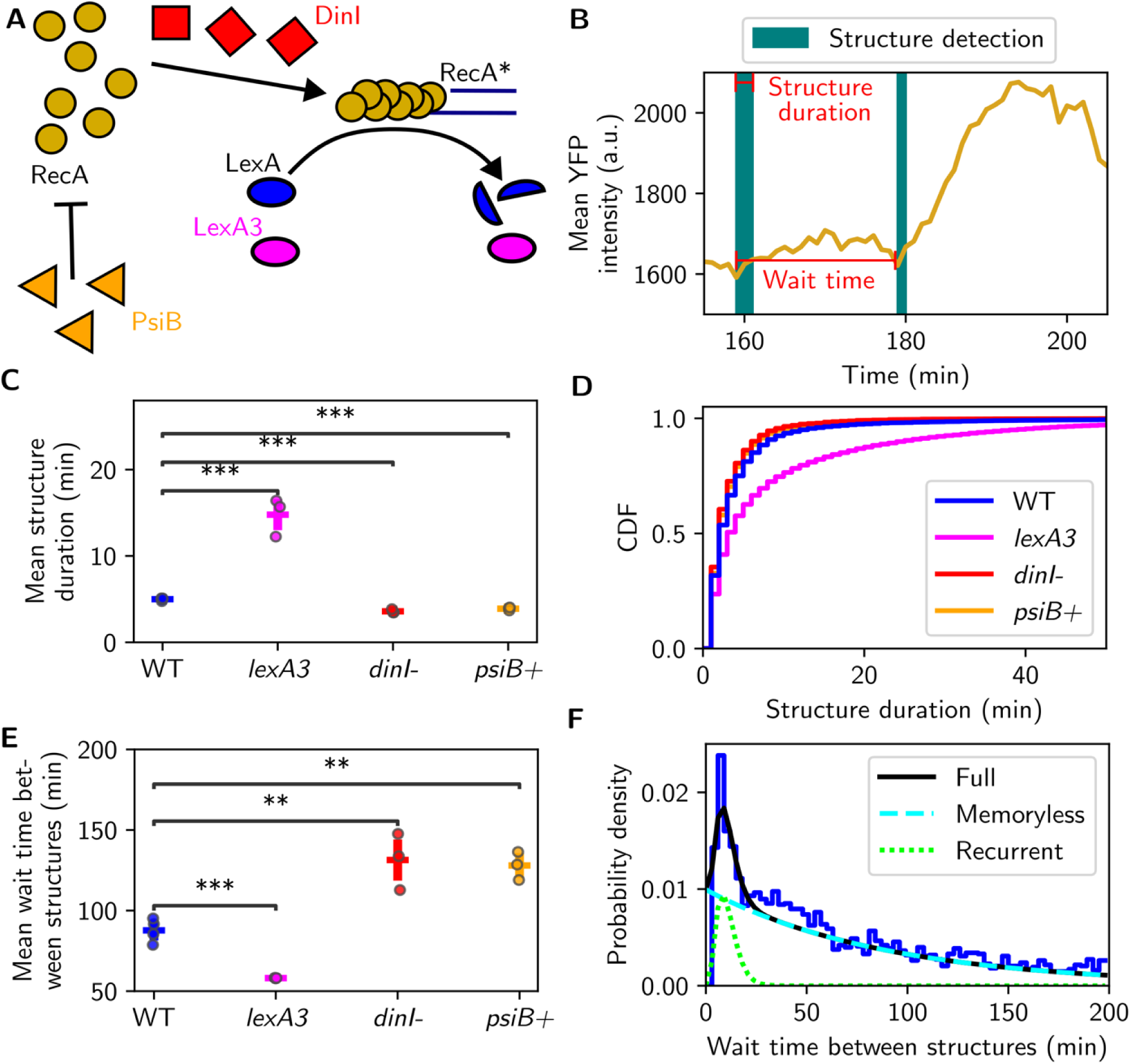
Analysis and genetic perturbation of RecA* structure kinetics. **(A)** Schematic of genetic perturbations. LexA3 is a non-cleavable LexA mutant that prevents SOS induction; DinI stabilizes RecA* structures; PsiB inhibits RecA* structure formation by binding cytoplasmic RecA. **(B)** Definitions of kinetic parameters. Structure duration is the time between first and last detection of a linked RecA* structure; wait time is the interval between successive structure onsets in single-cell time series. **(C)** Mean structure duration for each strain (mean ± s.e.m. across biological replicates; WT n=4; *lexA3, dinI-, psiB+* n=3). Statistical significance assessed using an unequal variance t-test. **(D)** Cumulative distribution functions of structure durations for each strain, pooled across biological replicates. **(E)** Mean wait time between structure onsets for each strain (mean ± s.e.m. across biological replicates; WT n=4; *lexA3, dinI-, psiB+* n=3). Statistical significance assessed using an unequal variance t-test. **(F)** Distribution of wait times from a representative replicate in WT strain (blue), with fitted probability density function (black). Contributions from memoryless and recurrent components are shown (dashed, turquoise and dotted, green lines, respectively).

For the wild type, the mean duration (Fig. 2B) of RecA* structures was 5 ± 1 min. The distribution of structure durations was heavily tailed, but only a minority of structures (35%) had lifetimes above 3 min (Fig. 2C,D). Destabilising RecA* structures via *dinI* deletion or reducing the free RecA pool via PsiB expression both reduced the average structure duration, as expected (Fig. 2C). In the *lexA3* mutant, not only were more structures observed in general, but the mean duration of structures increased 3-fold to 15 ± 2 min (Fig. 2C) and the tail in the structure lifetime distribution became far more pronounced (Fig. 2D). This suggests that long-lived RecA* structures correspond to repair intermediates whose resolution requires SOS induction. In the *lexA3* mutant, such lesions would not trigger expression of the required repair enzymes, leading to prolonged RecA* persistence. These RecA* structures frequently last longer than 30 min in the *lexA3* mutant. About half of observed RecA* structures remain short-lived in the *lexA3* mutant, indicating that these types of repair events can be completed without SOS activation. Thus, spontaneous RecA* structures in wild-type cells comprise at least two kinetic classes of repair intermediates: short-lived events that resolve independently of SOS induction, and rarer longer-lived events whose resolution is promoted by SOS-dependent repair functions.

### RecA* structure recurrence reveals memory in endogenous DNA repair dynamics

We next analysed the wait times (Fig. 2B) between successive RecA* structure formations, which had a mean value of 88 ± 4 min in unperturbed wild-type cells (Fig. 2E). For the *dinI-* and *psiB+* strains, the mean wait time increased relative to wild type (131 ± 10 min and 128 ± 5 min, respectively, Fig. 2E), suggesting that these perturbations delay the formation of RecA* structures. This is consistent with PsiB sequestering free RecA, which reduces the on-rate and therefore nucleation of structures. Similarly, the increased off-rate in the *dinI-*strain likely leads to a nucleation defect, where RecA* structures initially disassemble faster than they can grow. For the *lexA3* mutant, the mean wait time between RecA* structure formations of 58 ± 1 min was significantly lower than in the wild type (Fig. 2E). This suggests that the SOS response is important not only for resolving long-lived RecA* structures, but also for reducing the frequency of successive repair events.

To probe this further, we considered that DNA repair events could either be independent of each other (i.e. a memoryless process), or the onset of subsequent repair events could depend on previous instances of DNA damage (i.e. a recurrent process with memory). We found that a purely exponential fit, corresponding to a memoryless process of RecA* structure formation, was insufficient to explain the wait time distribution (Fig. 2F). We therefore fitted the distribution with a two-component model consisting of a long-timescale memoryless process and a short-timescale recurrence process. In wild-type cells, the mean wait time between memoryless events suggests that endogenous DNA damage events marked by RecA* structures occur on average every ∼100 min (∼2.2 cell generations). The mean wait time between recurrent events was shorter, at ∼12 min in wild-type cells. We interpret the short-timescale component as likely reflecting repeated attempts to process the same initial lesion. We confirmed by manual inspection of the raw fluorescence movies that RecA* structures disappeared between successive recurrent events, indicating that these are separate repair attempts. The wait time between memoryless events was substantially higher in *dinI-* and *psiB+* strains compared to the wild type (Supplementary Fig. 3 A), indicating that these perturbations delay the assembly of RecA* structures at independent damage events. In *lexA3* cells, the mean wait time between memoryless events was unchanged, suggesting that the rate of endogenous DNA damage events is unchanged. However, the proportion of recurrent events was higher in the *lexA3* mutant (Supplementary Fig. 3 B), suggesting that lack of SOS induction prevents damage resolution, leading to repeated repair attempts.

### RecA* structures are frequently associated with DNA replication forks in unperturbed cells

We next asked what the spatial and temporal properties of RecA* structures could reveal about the underlying causes of spontaneous SOS induction. Various endogenous processes generate stretches of ssDNA that can serve as a substrate for RecA loading. DNA replication is a major source of such ssDNA, as replication forks are vulnerable to stalling or collapse upon encountering DNA lesions, secondary structures, transcribing RNA polymerase or other DNA-bound protein obstacles (Kuzminov, 2001; Simmons et al., 2007; Merrikh et al., 2012; Gupta et al., 2013; Mei et al., 2021; Winterhalter et al., 2025). The frequency of spontaneous DNA breaks closely scales with the number of cell generations, supporting a replication-dependent mechanism (Shee et al., 2013). However, such damage can also arise independently of replication, for instance from reactive oxygen species or transcription-associated R-loops (Foster et al., 2015; Wimberly et al., 2013).

To test this directly, we imaged a strain co-expressing RecA-YFP and a beta-clamp fusion (DnaN-mCherry), which forms foci near active DNA replication forks (Mäkelä et al., 2021). The localisation patterns of RecA* structures and DnaN foci were very similar (Fig. 3A,B,C). Over the course of the cell cycle, both types of foci initially localised at mid-cell and moved towards quarter-cell positions as cells elongated (Fig. 3C), consistent with the known pattern of DNA replication in *E. coli* (Reyes-Lamothe et al., 2012). To quantify the association between RecA* structure formation and DNA replication, we computed the distance between the centroids of RecA* structures and the nearest DnaN foci, at the onset of each RecA* structure, in single cells (Fig. 3B). The distance distribution revealed that RecA* structures frequently appeared near DnaN foci (64% of onsets < 0.6 μm). Cells harbouring visible RecA* structures were significantly more likely to have DnaN foci, and vice versa (Fig. 3D,E). Taken together, a substantial fraction of spontaneous RecA* structures, and consequently spontaneous SOS induction events, arise in association with DNA replication during normal growth.

**Figure 3.**
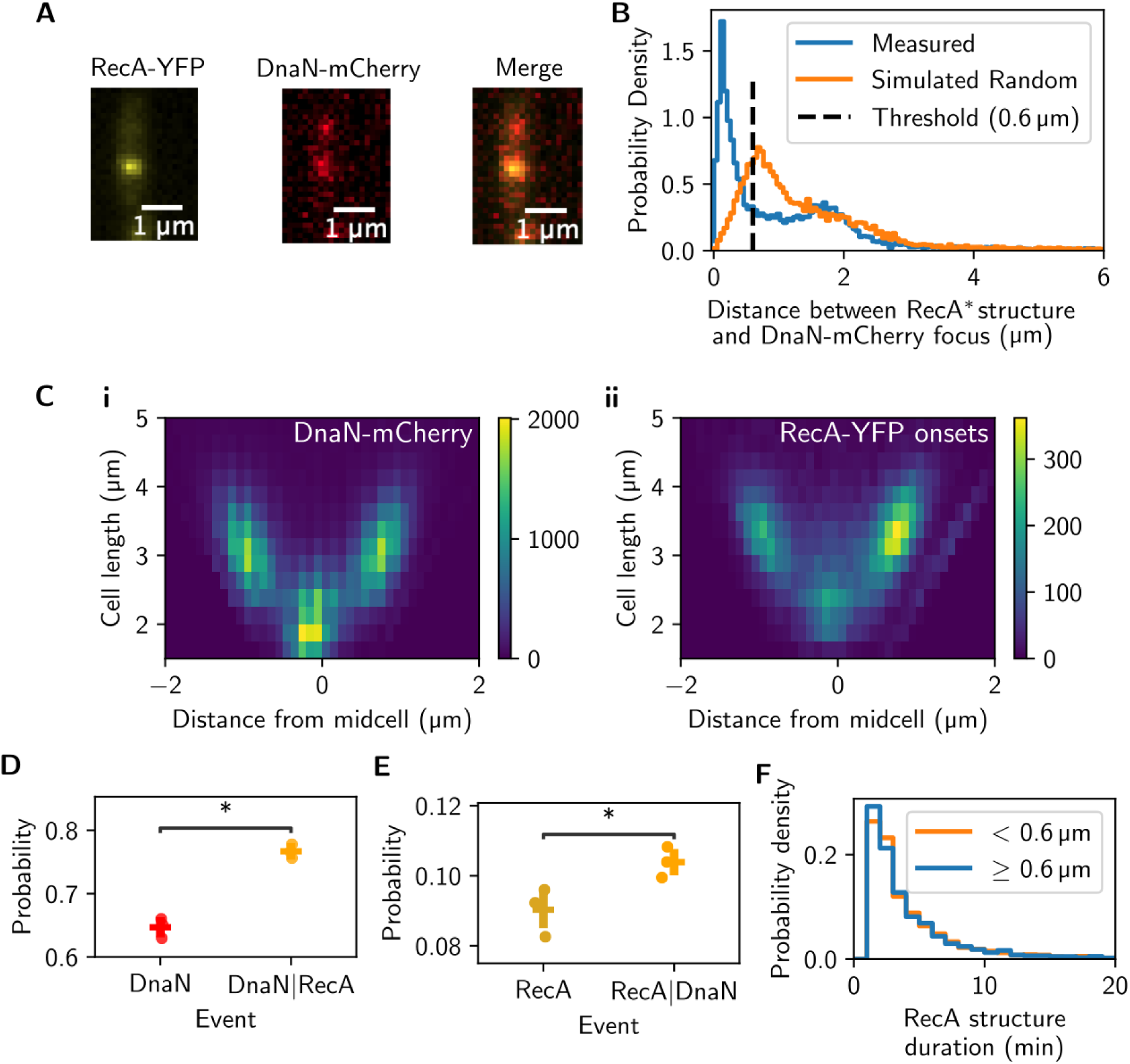
Association of RecA* structures with DNA replication forks. **(A)** Representative snapshot of single cell expressing RecA-YFP and DnaN–mCherry to mark replication foci, with merged image to assess co-localisation. **(B)** Distribution of distance between RecA* structure centroid and nearest DnaN–mCherry focus at the time of structure onset. Data were pooled across biological replicates (n=3). **(C)** (i) DnaN-mCherry foci localization and (ii) RecA-YFP localization at structure onset as a function of cell length. Data were pooled across biological replicates (n=3) and normalized by the maximum value in each vertical bin. **(D)** Probability of detecting a DnaN–mCherry focus either unconditionally or given the presence of a RecA* structure in the same cell (mean ± s.e.m. across biological replicates, n=3). **(E)** Probability of detecting a RecA* structure either unconditionally or given the presence of a DnaN–mCherry focus in the same cell (mean ± s.e.m. across biological replicates, n=3). Statistical significance in (D–E) was assessed using a paired t-test. **(F)** Distributions of RecA-YFP structure durations stratified by centroid distance from the nearest DnaN-mCherry focus (≤0.6 μm or >0.6 μm).

We examined whether the position of RecA* structures also influenced their kinetics. We reasoned that DNA breaks or ssDNA gaps occurring near a replication fork might be repaired more rapidly, as the recently duplicated sister chromosome copy provides an immediately available template for homologous recombination. In contrast, lesions arising elsewhere in the cell could require a more extensive homology search to locate an appropriate repair template (Lesterlin et al., 2014; Wiktor et al., 2021), potentially resulting in longer-lived RecA filaments. To test this hypothesis, we compared the lifetimes of RecA* structures that are proximal versus distant to DnaN foci, but found no significant differences (Fig. 3F). Therefore, although RecA* structures are often replication-associated, their subsequent dynamics, and consequently the magnitude of spontaneous SOS induction, are not measurably affected by fork proximity.

### Cumulative RecA* structure presence predicts spontaneous loss of growth and viability

We next examined the physiological consequences of spontaneous SOS induction, asking to what extent it contributes to the resolution of endogenous DNA damage that would otherwise compromise cell viability. Using the mother machine device, cell fates were quantified by tracking the growth behaviour of individual mother cells over time (Wang et al., 2010). We identified four main categories of cell fate. Cell death was defined by a sudden loss of cytoplasmic fluorescence, indicative of lysis. A second class of cells exhibited sustained growth arrest, while a third underwent filamentation, leading in a small number of cells to excessive filamentation and escape from the growth channels. For the purposes of our analysis, these three outcomes (lysis, growth arrest, and filamentation loss) were collectively defined as an “inactive” state. The remaining cells were classified as actively growing. The wild type, *dinI-*, and *psiB+* strains exhibited comparable percentages of actively growing cells after 520 min of imaging (chosen as a fixed measurement window), whereas survival of *lexA3* mutant cells was significantly lower (Fig. 4A,B). This confirms that while alterations in RecA filament nucleation or stability do not impair survival of endogenous DNA damage (Renzette et al., 2007), inhibiting downstream SOS gene induction strongly increases spontaneous cell death rates (Wang et al., 2010).

**Figure 4.**
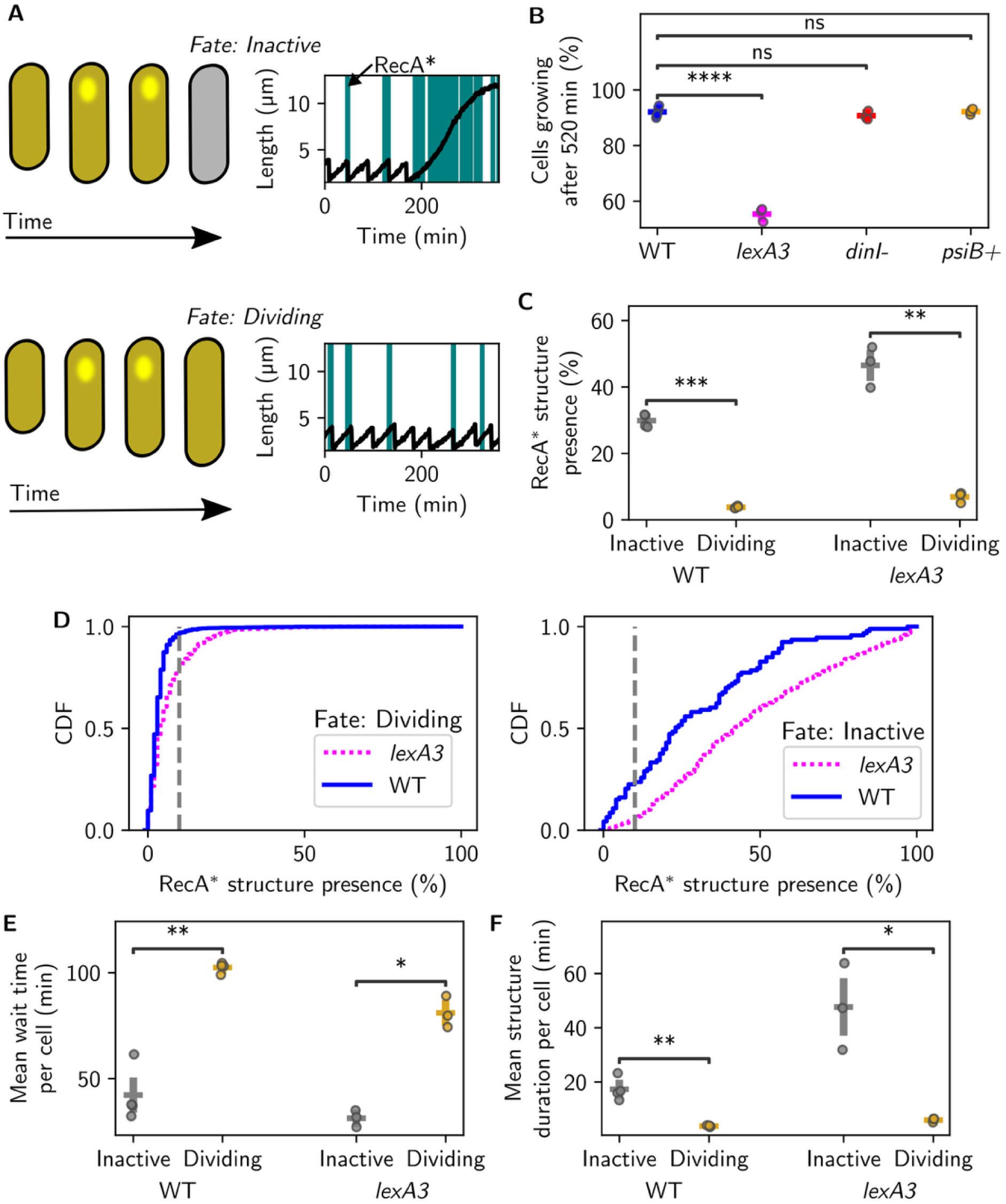
Cumulative RecA* structure presence predicts cell fate. **(A)** Schematic of cell fate classification as ‘dividing’ (actively growing) or ‘inactive’ (including lysis, growth arrest, or filamentation escape) and definition of cumulative RecA* structure presence as the percentage of the active growth period for which a RecA* structure is detected, computed from single-cell time series. **(B)** Fraction of the initial population remaining actively growing at 520 min (mean ± s.e.m. across biological replicates; WT n=4; *lexA3, dinI-, psiB+* n=3). Statistical significance assessed using an unequal variance t-test. **(C)** Average percentage of time spent with RecA* structure present for each cell, stratified by fate at 520 min (mean ± s.e.m. across biological replicates; WT n=4; *lexA3* n=3). **(D)** Cumulative distributions of RecA* structure presence in single cells, stratified by fate at 520 min. Data pooled across biological replicates for each strain (WT n=4; *lexA3* n=3). The grey dashed line represents the 10% threshold used to differentiate between cells with high and low structure presence. **(E)** Mean RecA* structure duration stratified by cell fate (mean ± s.e.m. across biological replicates; WT n=4; *lexA3* n=3). **(F)** Mean wait time between RecA* structure onsets, stratified by cell fate (mean ± s.e.m. across biological replicates; WT n=4; *lexA3* n=3). Statistical significance in (C,E,F) was assessed using a paired t-test.

We tested whether the presence of RecA* structures could predict individual cell fate. Cells were classified as dividing or inactive based on their status at 520 min, and the fraction of time each cell exhibited detectable RecA* structures was quantified (Fig. 4C). For cells that became inactive, this measure was calculated only over the period preceding loss of activity (Fig. 4A,C). This metric of cumulative RecA* structure presence captures both the frequency and duration of structures while normalizing for observation time. Cells that became inactive spent a significantly larger fraction of their lifespan with RecA* structures compared to dividing cells, indicating that persistent or frequent RecA* structures strongly predict spontaneous loss of activity during unperturbed growth. To further quantify this relationship, we examined the full distribution of RecA* structure presence across individual cells. Cumulative RecA* presence was broadly distributed, ranging from near 0% to almost 100% of the observation time per cell (Fig. 4D). When grouped by fate, these distributions diverged markedly: 97% of dividing cells exhibited low RecA* presence (<10%; grey dashed line threshold in Fig. 4D), whereas only ∼25% of cells that became inactive also showed low RecA* presence (<10%) (Fig. 4D). Thus, most cells (75%) undergoing lysis, growth arrest, or filamentation exhibited elevated RecA* structure presence, identifying endogenous DNA damage and repair activity as a prominent feature of spontaneous departures from normal growth.

### Spontaneous loss of growth is associated with repeated DNA damage and repair failure

The increased presence of RecA* structures in the inactivity-fated cells could arise from two, not mutually exclusive, scenarios. In the first, a single endogenous DNA damage event proves exceptionally difficult to repair, resulting in prolonged RecA* structure persistence and ultimately cell death. In the second, a cell experiences more frequent DNA damage events and correspondingly shorter interval between RecA* structures, increasing the cumulative likelihood of eventual death. To distinguish between these possibilities, we analysed our single-cell data from two perspectives. In the first approach, we stratified RecA* structures by their duration, and calculated for each structure event the rate of a cell transitioning from active to inactive fate before the next structure event. This rate was quantified as a function of the structure’s duration. However, both the WT and *lexA3* mutant showed no significant association between RecA* structure duration and rate of transition to inactivity (Supplementary Fig. 4). Thus, a single, long-lived RecA* structure was not by itself predictive of imminent cell lysis or growth arrest.

In the second approach, we examined the mean duration and mean wait time of all detected RecA* structures within a cell stratified by its fate at 520 min (Fig. 4E,F). In cells that eventually became inactive, structure durations were significantly longer and wait times significantly shorter than in the dividing population. Therefore, DNA repair events are both more frequent and more persistent over the lifetime of cells that later become inactive. Consistent with our earlier observation that RecA* structure formation exhibits memory, with events often occurring in rapid succession (Fig. 2E), these results point to an increased overall burden of DNA damage and repair in inactivity-fated cells. Together, this supports a cumulative-damage model (Yang et al., 2023), in which repeated chronic DNA damage and repair activity, rather than a single catastrophic event, is associated with most spontaneous cell death or growth arrest events during unperturbed growth.

### Endogenous DNA damage precedes physiological decline

Having established that cells fated for inactivity experience a higher DNA damage burden, we asked when this fate divergence first becomes detectable. To do so, we aligned single-cell traces by the time of lysis, or arrest, and computed the mean number of RecA* structures per cell as a function of time before each fate. Strikingly, the frequency of RecA* structures began to rise approximately 4 hours before death or arrest, closely followed by increased SOS expression (Fig. 5). This shows that elevated RecA* activity is a very early indicator of eventual cell death, appearing around 5-6 cell generations in advance. We next studied the temporal order of events leading to spontaneous death. We found that RecA* structures appeared significantly earlier than any detectable change in cell growth rate or morphology (Fig. 5). In particular, cell growth rate began to decline only ∼100 min prior to death or arrest, substantially later than the increased RecA* activity. Hence, elevated RecA* activity is an early detectable event in the trajectory toward spontaneous cell death, rather than a late consequence of overt physiological decline, appearing approximately 2 hours before major physiological perturbations become apparent.

**Figure 5.**
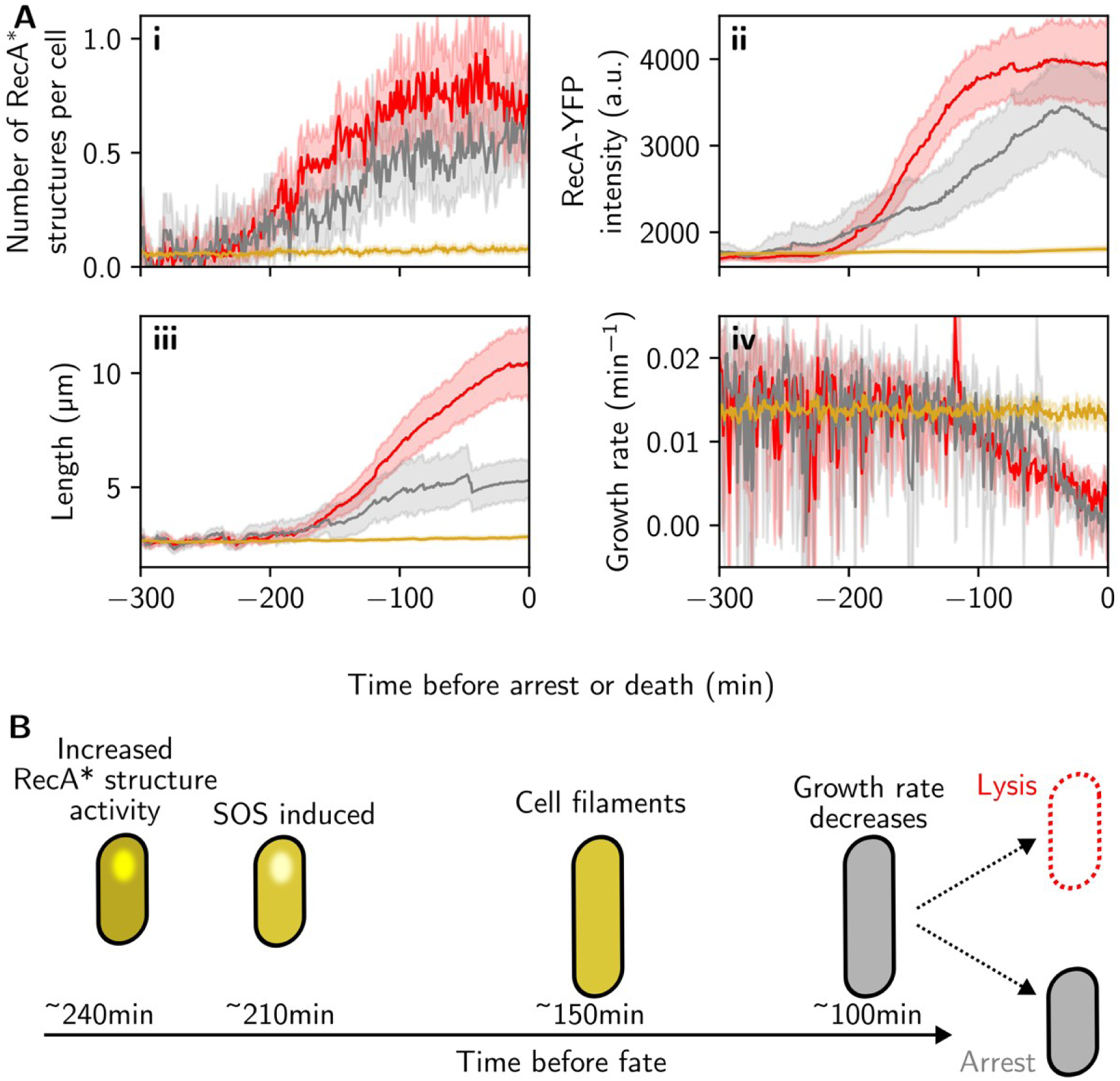
Endogenous DNA damage events are early indicators of unprovoked cell death or growth arrest. **(A)** Single-cell traces were aligned in time relative to the moment of lysis or growth arrest (set to time = 0). Time-aligned population averages for (i) number of RecA* structures per cell, (ii) RecA-YFP intensity, (iii) cell length, and (iv) growth rate in cells that lyse (red), arrest (grey), or continue dividing. For dividing cells, traces were aligned to an arbitrary reference timepoint. Data show mean ± s.e.m. (solid lines and shaded areas) over cell populations pooled over biological replicates, giving n=71 for lysing cells, n=52 for arresting cells, and n=1313 dividing cells. **(B)** Summary of observed chronology of cell phenotypes prior to lysis or growth arrest.

## Discussion

The SOS response is one of the best-characterized bacterial stress-response pathways, with decades of work establishing the central functions of RecA, LexA, and SOS-dependent DNA repair. Most mechanistic studies, however, have focused on population-level responses to acute exogenous damage. Single-cell studies have shown that SOS induction also occurs spontaneously during unperturbed growth, but why these activation events vary so strongly between cells has remained unclear. Our results place the major source of variability upstream of gene expression, at the level of RecA* structure formation and resolution, in agreement with theoretical predictions (Galbusera et al., 2025). Although fluctuations in LexA binding, transcription, translation, and protein turnover may contribute to SOS heterogeneity, our data show that spontaneous SOS pulses are finely graded according to the lifetime and size of RecA* structures. In this way, graded SOS induction can match the cellular response to the difficulty of endogenous DNA damage processing. Consistent with this view, quantitative changes in RecA abundance have recently been shown to shift the balance between alternative DNA repair activities, suggesting that graded levels of SOS induction may control not only overall repair capacity, but also repair pathway choices (Fleurier and Matic, 2026). While short-lived repair intermediates can often be resolved by basal repair capacity, long-lived or recurrent intermediates depend on SOS-induced repair functions. Similar logic applies to eukaryotic DNA damage responses, where ATM/ATR and p53 dynamics encode the extent and persistence of DNA damage to tune cell fate decisions between repair, arrest, and death (Ciccia and Elledge, 2010; Paek et al., 2016). Thus, although SOS and eukaryotic DNA damage-response pathways are molecularly distinct, both systems illustrate how cells convert damage-processing dynamics into graded cell-state responses.

An apparent conundrum from our data is why short-lived RecA* structures induce any SOS response if many of these repair intermediates can be resolved without SOS-dependent gene expression. This question is particularly relevant because bacteria in natural environments are unlikely to experience only the extreme acute stresses often used experimentally (Andersson and Hughes, 2014; Imlay, 2019); instead, they frequently encounter fluctuating, moderate, or sublethal stresses for which transient preparedness may be advantageous. Such anticipatory stress responses are conceptually related to bet-hedging, where heterogeneous or transient activation of costly protective pathways can improve survival under uncertain future conditions (Ackermann, 2015; Bayramoglu et al., 2017; Lagage and Uphoff, 2020). One possibility, therefore, is that cells use each endogenous DNA damage event as a warning signal of increased future risk. Even when the initiating lesion can be resolved at basal repair capacity, transient SOS induction may raise the abundance of repair and damage-tolerance proteins, thereby facilitating the resolution of subsequent lesions or preventing partially repaired intermediates from progressing into more severe damage. Consistent with this idea, *lexA3* cells can still resolve many short-lived RecA* structures, but show a reduced mean wait time between successive structures, suggesting that the absence of SOS induction increases the probability of recurrent repair events. In this view, spontaneous SOS activation is not only a response to the current lesion, but also a short-term protective state that anticipates further damage.

Indeed, we find that endogenous DNA damage is not an incidental stress during optimal growth, but a major route by which individual cells depart from normal proliferation. This happens even in cells with fully operative repair systems, and has been linked to the presence of cell subpopulations experiencing endogenous physiological stresses and increased mutation rates (Gonzalez et al., 2008; Woo et al., 2018). Cells rely on many essential systems whose failure could, in principle, cause death, including metabolism, envelope maintenance, protein homeostasis, chromosome segregation, and cell division. Yet in our experiments, most cells that later lysed, arrested, or filamented had previously experienced increased RecA* structure presence. Importantly, these fate trajectories were associated with persistent and recurrent repair intermediates rather than isolated catastrophic events, and became detectable several generations before changes in growth rate or morphology. Cumulative failure of DNA maintenance may therefore represent a major source of hidden physiological heterogeneity in growing bacterial populations.

## Acknowledgements

Members of the Uphoff, Baker, and Maini groups are thanked for useful comments and discussions. This research was funded by a Wellcome Trust Sir Henry Dale Fellowship (206159/Z/17/Z) and a Research Prize Fellowship by the Lister Institute of Preventative Medicine to SU. JJP was supported by St. John’s College Oxford and an Oxford Interdisciplinary Bioscience Doctoral Training Partnership studentship by the Biotechnology and Biological Sciences Research Council (BB/T008784/1). AC acknowledges a Research and Training Grant by the Federation of European Microbiological Societies (FEMS). REB acknowledges support of a grant from the Simons Foundation (MP-SIP-00001828).

For the purpose of open access, the authors applied a CC BY public copyright license to any author accepted manuscript arising from this submission.

## Materials and methods

### Bacterial strains

All bacterial strains are derivatives of E. coli AB1157. Alleles encoding fluorescent protein fusions or gene deletions were incorporated into this background strain using P1 phage transduction and selection via the appropriate antibiotic resistance marker (kanamycin, tetracycline). The *recA-recA-syfp2 kan* allele was kindly provided by Johan Elf’s lab (Wiktor et al., 2021). To aid cell segmentation, strains constitutively expressed cytoplasmic fluorescent proteins mKate2 (or msCFP3) from a constitutive promoter Prna1. Gene deletion mutants containing a kanamycin resistance gene (*frt-kan-frt*) were obtained from the Keio collection (Baba et al., 2006). The regulator of flagella synthesis, FlhD, was deleted to render *E. coli* non-motile for confinement within microfluidic channels (Uphoff et al., 2016). To combine multiple alleles using P1 transduction with kanamycin selection, the kanamycin resistance gene of the intermediate strain was first removed by transformation with plasmid pCP20. Growth at 37 °C induced recombinase expression to loop out the frt-flanked resistance gene and cured the temperature-sensitive plasmid. Insertions and deletions were verified by antibiotic selection, PCR and gel electrophoresis, and microscopy (for fluorescent proteins). PsiB was expressed from a pSC101 plasmid under a Plac promoter.

**Table 1:**
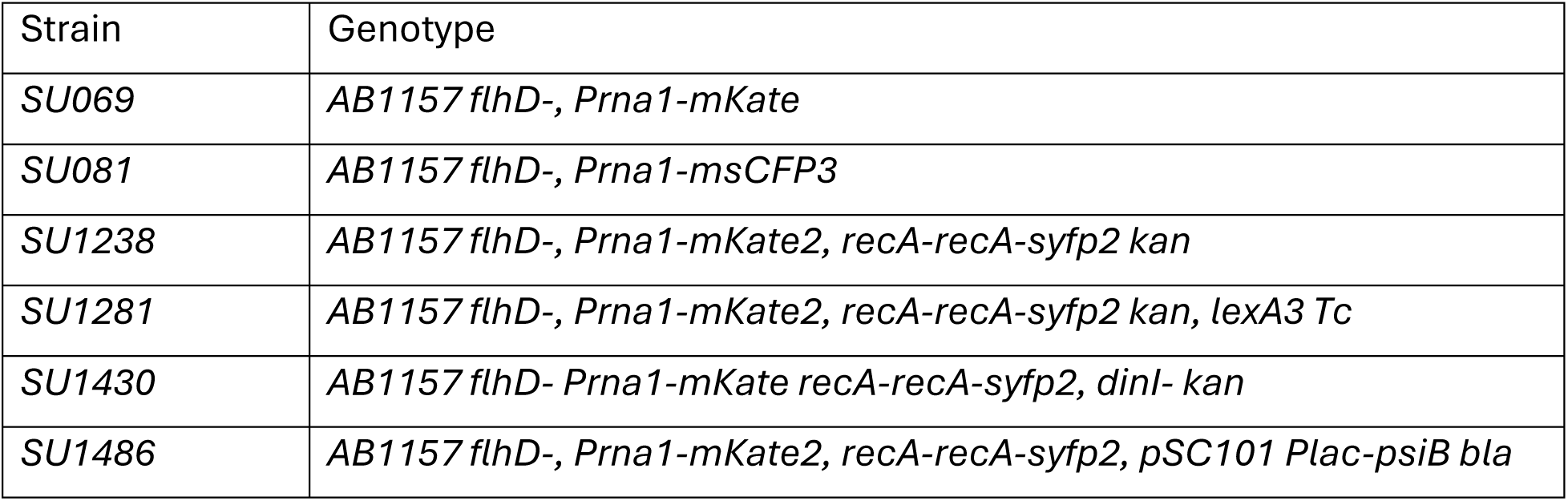
List of *E. coli* strains.

### Bacterial culture

*E. coli* strains were stored at -70°C as frozen glycerol stocks, and streaked on LB agarose plates with appropriate antibiotic. For each experiment, a single colony was picked and grown in LB media at 37 °C with shaking at 200 rpm. After 4 h, cultures were diluted 1:2000 in 4 ml M9 minimal media and grown for 18-24 hours over night. The next day, culture was diluted 1:50 in M9 minimal media with 0.85 mg ml^−1^ pluronic acid F127 (Sigma) and grown for 3 h, then centrifuged for 3 min at 6000 rpm and resuspended in 30 μl of supernatant before loading into mother machine devices. M9 minimal media contained 100 μM CaCl_2_, 2 mM MgSO^4^, M9 salts (Sigma-Aldrich), MEM amino acids (Gibco), 100 mg ml^−1^ L-proline (BioChemica), 5 × 10^−5^ % Thiamine (BioChemica), 0.2% Glucose.

### Preparation of microfluidic devices

Mother machine microfluidic experiments were performed as previously described (Wang et al., 2010; Uphoff et al., 2016). Mother machine devices were prepared from Polydimethylsiloxane (PDMS) monomer mixed with curing agent in ratio 1:10 (Dow Corning Sylgard 184, Farnell electronics). The mixture was poured onto a silicon wafer (ConScience) and air bubbles were removed under vacuum, then cured for 2 h at 65 °C, rinsed with 100% ethanol and dried using nitrogen gas. Media inlet and outlet holes were inserted into devices using a 0.75 mm biopsy punch. Coverslips (no 1 thickness) were cleaned by sonication in acetone for 20 min, rinsed with MilliQ and sonicated in 100% isopropanol for 20 min, then dried using nitrogen gas. The coverslip and PDMS device were treated with air plasma (Plasma Etch PE-50) before bonding for 30 min at 95 °C.

Concentrated cell suspension (prepared as described above) was inserted into device inlets and outlets via pipetting, before centrifuging the device in a custom-built holder at 5000 rpm to load cells into the mother machine trenches. Syringes (Terumo or BD plastipak) were filled with M9 minimal media containing 0.85 mg ml^−1^ pluronic acid F127 to reduce cell adhesion. If indicated, media contained additionally 1 ng ml^−1^ ciprofloxacin (Sigma-Aldrich) for DNA damage treatment. Syringes were attached to a syringe pump (ALADDIN-220, World Precision Instruments) connected through Tygon Microbore silicon tubing (VWR ND100-80/0.508*1.524) to the device. The device outlet was connected to a waste container sealed with Parafilm. To flush out excess cells from inlet and outlet channels, the flow rate was set to 2.5 ml/h for 15 min, then reduced to 0.5 ml for the remainder of the experiment (typically up to 18 h). Cells were allowed to grow in the device for at least 2 h prior to imaging. To add ciprofloxacin treatment, flow was temporarily paused and a syringe containing the antibiotic was swapped into the pump, with flow then restarted.

### Fluorescence microscopy

Cells in microfluidic devices were imaged by time-lapse fluorescence microscopy with a Nikon Ti Eclipse inverted microscope. The setup had a perfect focus system, oil immersion objective (100x, NA 1.4), motorised stage, sCMOS camera (Hamamatsu Flash 4) and LED excitation source (Lumencore Spectra X). LED intensities and exposure times were set to 50% and 300 ms for RecA-SYFP2 or SYTOX Green, 50% and 100 ms for mKate2 or mCherry-DnaN, 7.5% and 100 ms for msCFP3. The microscope’s temperature chamber (Okolabs) was maintained at 37 °C. Time-lapse movies were recorded at 1 min time interval for 18 positions. Each position had 20 growth trenches, producing up to 360 mother cell traces per experiment. The microscope was operated using NIS-Elements software (Nikon).

### Mother machine data processing

Movies were analysed using a combination of software and code: Fiji (Schindelin et al., 2012), Bacteria in Mother Machine Analyzer (BACMMAN) (Ollion et al., 2019), and a custom-built Python pipeline inspired by previous work (Choudhary et al., 2023), Visual Studio Code (Microsoft).

In BACMMAN, images were processed using the following functions:

- RemoveHotPixels: Removes single pixels of substantially higher intensity than surroundings in space and time.
- RemoveStripesSignalExclusion: Removes banding noise.
- SaturateHistogramHyperfluoBacteria: Detects bacteria of very high fluorescence intensity relative to other bacteria and saturates the pixels corresponding to those bacteria.
- AutoRotationXY: Rotates images to align microchannels with the *y*-axis.
- CropMicrochannelsFluo2D: Crops images around the microchannels.
- TypeConverter: Lowers bit depth of pre-processed images to save memory.
- MicrochannelFluo2D: Detects mother machine trenches.
- BacteriaClosedMicrochannelTrackerLocalCorrections: Bacterial segmentation using marker fluorescence.
- NestedSpotTracker: For RecA^∗^ structure or DnaN foci detection.

Structure tracking was performed using BACMMAN TrackMate (Tinevez et al., 2017).

BACMMAN outputs were exported as .csv files for further analysis and data visualization. The following data were recorded:

- Mean fluorophore intensity: Mean pixel intensities within a segmented bacterium.
- Mean structure intensity: Mean pixel intensities within a segmented structure.
- Cell area: Total area within a bacterium’s segmented contour.
- Cell length: Maximum distance between two points of bacterium’s segmented contour.
- Number of structures per cell: Number of segmented structures within a bacterium’s segmented contour.
- Structure area: Total area within a foci/structure’s segmented contour

We calculated the following quantities using Python:

- Growth rate: We smoothed Cell Length time series by applying the pandas `.rolling().mean()` function with windowsize=5. For the smoothed length timeseries *L̃_ι_* at each timepoint *i*, we calculated the logarithm of the ratio between adjacent lengths ln(*L̃_i_*/*L̃_i_*_−1_). To remove cell division events, values of ln(*L̃_i_*/*L̃_i_*_−1_) < ln(0.9) were discarded. Given a frame interval Δ*t*, the growth rate was calculated as ln(*L̃_i_*/*L̃_i_*_−1_)(Δ*t*)^−1^.
- Growth arrest: We used pandas `.rolling().std()` function with windowsize=40 to obtain the rolling standard deviation of the cell length, *s_L_*. A threshold of *s_L_* < 0.2 was applied to identify periods of arrested growth and assigned corresponding frames a ‘growth score’ *S_i_* of 0. Since we expect growth arrest to be sustained, we applied a second threshold, requiring that over a window of 12 frames at frame *j*, the 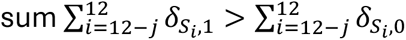, where *δ*_k,l_ is the Kronecker delta, which is 1 if *k* = *l* and 0 otherwise. This additional filter incorporates memory into the detection of cell growth arrest; if a cell has been detected as growing over all previous frames within the 12-frame window, the threshold to be detected as arrested is higher.
- Lysis or filamentation escape were detected based on loss of fluorescence segmentation marker signal. Filamentation escape was only concluded if the cell length prior to fluorescence loss was greater than 20 μm.
- RecA^∗^ structure duration and wait time: The time difference between the last and first frame of a tracked RecA* structure, and the interval between two separate RecA* structures within the same cell.
- Number of RecA* structures in each frame divided by cell length.
- Total RecA* structure intensity within segmented area of a structure and corrected for background intensity.
- RecA* structure presence: Fraction of time each cell exhibited detectable RecA* structures prior to cell death or loss of activity.
- Probability of RecA^∗^ structure P(RecA): Sum of frames where at least one RecA^∗^ structure was present divided by the number of frames observed.
- Conditional probability of RecA^∗^ structure given DnaN focus is present in cell P(RecA|DnaN): Sum of frames where at least one DnaN focus and at least one RecA^∗^ structure were present simultaneously in the cell divided by the number of frames for which only DnaN foci were present.
- Probability of DnaN foci P(DnaN): Sum of frames where at least one DnaN foci was present divided by the number of frames observed.
- Conditional probability of DnaN foci given RecA^∗^ structure is present in cell P(DnaN|RecA): Sum of frames where at least one DnaN focus and at least one RecA^∗^ structure were present simultaneously in the cell divided by the number of frames for which only RecA^∗^ structures were present.
- Distance between RecA^∗^ structure and DnaN focus at onset of structure.
- Distribution of RecA* structure wait time: We used scipy.stats.fit to fit the distribution of wait time between RecA^∗^ structures with the probability density function 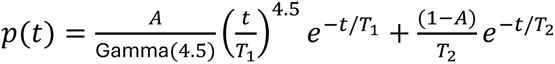. Here, *A* corresponds to the proportion of recurrent structure events (with memory), 4.5 · *T*_1_ corresponds to the mean lifetime of recurrent events, and *T*_2_ corresponds to the mean lifetime of memoryless events. Gamma corresponds to the mathematical gamma function. To reduce the number of free parameters associated with the Gamma distribution, we constrained the shape parameter to 4.5 as shown above. Similar fits can be obtained by fixing the parameter *T*_1_ instead. Fit parameters were obtained from the mean of 300 fitting attempts for *T*_2_ and *A*.
- Rate from dividing to inactive population before next structure as a function of structure duration: Mean over transition rates associated with all structures with the same duration. The transition rate calculation for each structure depends on the cell fate classification or presence of another structure following that structure:
  - For all structures followed immediately by an ‘inactive’ fate declaration, the transition rate is the reciprocal of the number of frames between structure disappearance and the fate declaration.
  - For all structures followed by another structure or, no structure but a ‘dividing’ fate declaration at the final timepoint, the transition rate is zero.

### Simulated distribution of distance between random localisations of RecA* structures and DnaN foci

Idealised RecA* structure centroid positions were generated according to a uniform probability distribution within a cylindrical region with hemispherical end caps of radius 0.34 μm. Using each measured DnaN foci centroid position, the distance between each RecA* structure and simulated foci was calculated.

### SOS response mathematical model

We generated a system of ordinary differential equations (ODEs) to model the dynamics of SOS response gene regulation, based on established models (Ronen et al., 2002; Galbusera et al. 2025):

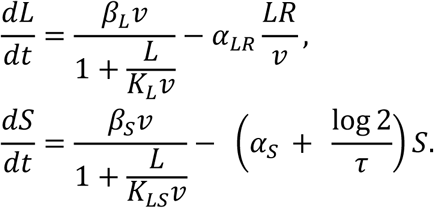

Here, *L*, *S* represent the number of LexA molecules and RecA-YFP mean fluorescence intensity, respectively. *v* represents the cell volume and *R* represents the number of RecA^∗^ structures present in the cell. *β_L_*, *β_S_*, *K_L_*, *K_LS_*, *α_LR_*, *α_S_*, *τ* are positive constants (Table 2), chosen for qualitative agreement between experimental observations and model outputs, and similar to previous ODE models of the SOS response (Krishna et al., 2007; Ronen et al., 2002). For *L*, the first term represents production of *L* and the second term represents LexA degradation when a RecA^∗^ structure is present. For *S*, the first term represents production of *S* and the second term represents degradation and photobleaching of *S*.

**Table 2:** Parameters of SOS response model.

| Parameter | Value | Interpretation |
| --- | --- | --- |
| $K_L$ | 210 nM | Dissociation constant for LexA-*PlexA*. |
| $K_{LS}$ | 80 nM | Dissociation constant for LexA-*PrecA*. |
| $\alpha_{LR}$ | $4.5 \text{ min}^{-1} \mu\text{m}^3$ | Degradation rate of LexA (per unit volume) due to presence of a RecA* structure. |
| $\alpha_S$ | $0.01 \text{ min}^{-1}$ | Degradation rate of RecA-YFP. This rate implicitly contains the conversion factor from concentration to fluorescence intensity. |
| $\tau$ | 49.2 min | Generation time. |
| $\beta_L$ | 1330 nM | Maximum production rate of $L$ when repressor concentration is zero. Chosen using $K_L$ and $\alpha_{LR}$ such that the steady-state concentration of $L$ is fixed at 1300 nM. |
| $\beta_S$ | $247 \text{ a.u. min}^{-1}$ | Production rate of $S$ in absence of $L$. Chosen such that the steady state mean intensity is close to the mean, background-subtracted, uninduced RecA-YFP intensity observed experimentally. |
| $k_1$ | $0.01 \text{ min}^{-1} \mu\text{m}^{-3}$ | Onset rate of RecA* structures per unit volume. |
| $k_2$ | $0.2 \text{ min}^{-1}$ | Rate of disappearance of RecA* structures. |
| $v_0$ | $0.75 \mu\text{m}^3$ | Cell volume at birth. |

We assume equilibria in promoter binding-unbinding according to the current concentration of *L*. This assumption has been shown to be an oversimplification in achieving quantitative agreement with experimental data (Culyba et al. 2018; Galbusera et al. 2025). However, this assumption is still sufficient to qualitatively capture the effect of RecA^∗^ structure kinetics, which depend only on the modulation of LexA degradation rate due to an input control parameter (in this case, *R*).

Exponential cell growth was described by a simplified adder model with doubling time *τ* (Wallden et al., 2016):

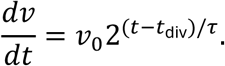

For each simulated cell, *t*_div_ was drawn from a uniform a distribution U[−*τ*, 0] to produce an initial condition for *v*. The cell volume was simulated using the Scipy function solve_ivp (Virtanen et al., 2020). When the cell volume reached *v* = 2*v*_0_, the volume was set to *v*_0_, and *t*_div_ set to the current time.

For each cell, a timeseries of RecA* structure events *R* was modelled using a stochastic birth-death process,

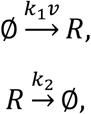

with positive constants *k*_1_ and *k*_2_, using the Tau Leap algorithm with timestep 0.005 min (Gillespie, 2001), implemented in Python. The birth-death reactions represent the spontaneous production and removal of RecA* structures with rates *k*_1_*v* and *k*_2_, respectively. At cell division events *v* = 2*v*_0_, any RecA* structures present were retained in the current cell.

Using the timeseries for *v* and *R*, timeseries for LexA concentration and RecA-YFP mean intensity were simulated using Scipy’s solve_ivp and their respective ODEs. *L* was halved upon cell division events (when *v* = 2*v*_0_). This ensured that LexA concentration remained constant immediately before and after cell division.

2000 simulated cells were obtained, each simulated for 700 min. The first 150 min, where the mean intensity of RecA-YFP over all cells was not yet at steady state, were discarded to remove the dependence of timeseries on an arbitrary choice of initial condition. SOS response curves were generated from the values of *L* and *S* following the onset of *R* events, then averaged over all events of a given structure duration from 2000 simulated traces of 600 min duration.

